# MtExpress, a Comprehensive and Curated RNAseq-based Gene Expression Atlas for the Model Legume *Medicago truncatula*

**DOI:** 10.1101/2021.05.27.445921

**Authors:** Sébastien Carrère, Jérôme Verdier, Pascal Gamas

## Abstract

Although RNA sequencing has been becoming the main transcriptomic approach in the model legume *Medicago truncatula*, there is currently no genome-wide gene expression atlas covering the whole set of RNAseq data published for this species. Nowadays, such tool is highly valuable to provide a global view of gene expression in a wide range of conditions and tissues/organs.

Here, we present MtExpress, a gene expression atlas that compiles an exhaustive set of published *M. truncatula* RNAseq data (https://medicago.toulouse.inrae.fr/MtExpress). MtExpress makes use of recent releases of *M. truncatula* genome sequence and annotation, as well as up-to-date tools to perform mapping, quality control, statistical analysis and normalization of RNAseq data. MtExpress combines semi-automated pipelines with manual re-labelling and organization of samples, to produce an attractive and user-friendly interface, fully integrated with other available Medicago genomic resources. Importantly, MtExpress is highly flexible, in terms of both queries, e.g. allowing searches with gene names and orthologous gene IDs from Arabidopsis and other legume species, and outputs, to customize visualization and redirect gene study to relevant Medicago webservers.

Thanks to its semi-automated pipeline, MtExpress will be frequently updated to follow the rapid pace of *M. truncatula* RNAseq data publications, as well as the constant improvement of genome annotation.

## Introduction

Legumes represent a plant family of major importance for food, feed and the agroecosystem, thanks to their nutritional qualities and their ability to grow efficiently in the absence of nitrogen fertilizers due to their endosymbioses with soil microorganisms, notably with nitrogen-fixing soil bacteria (rhizobia). Indeed, legumes are becoming a keystone in tomorrow’s agriculture to ensure food security, sustainable practices and mitigation of greenhouse effects. *Medicago truncatula* was proposed as a model legume about thirty years ago, and since then it has been widely used to study a wide range of biological issues relevant for plant biology and legume crop improvement. Its model legume status has led to the development of important resources for genetics, reverse genetics, structural and functional genomics (Kang et al. 2016). This notably included a gene expression atlas (MtGEA; https://mtgea.noble.org/v3) based upon microarray technology developed by Affymetrix and representing 33,229 genes from the genome version 3 (Kang et al. 2016). This atlas integrated 270 different samples (not considering biological replicates) in its last release and has been extensively used by the Medicago community. Unfortunately, MtGEA website was not maintained and no longer responds to inquiries.

In the past years, RNA sequencing has become increasingly popular to study gene expression at the plant, organ and tissue levels. The advantages of this approach are its sensitivity and its flexibility to exploit new genome releases and annotations by the possibility of (re)mapping sequencing reads, as well as to discover transcript isoforms or novel transcripts such as long noncoding RNAs. Two databases (db) give access to some *M. truncatula* RNAseq data, the first one as part of a generic db developed by EBI on many species (https://www.ebi.ac.uk/gxa/plant/experiments), and the second one as part of a specific study dedicated to nutrient- and nodulation- responsive peptides (http://mtsspdb.noble.org/atlas-internal/3880/transcript/profile/0) (de Bang et al. 2017). However, these db only show a small selection of publicly available RNAseq data. Here we present MtExpress, a gene expression atlas based upon *M. truncatula* RNAseq data supported by peer-reviewed publications (https://medicago.toulouse.inrae.fr/MtExpress). We ensured that MtExpress was a comprehensive, reliable, user-friendly, interconnected and upgradeable database, by combination of automated analyses and manual curation.

## Results

### Retrieval, quality checking and organization of RNAseq data

To build MtExpress, RNAseq data (i.e. sequenced reads, fastq) generated from Medicago were downloaded from the Sequence Read Archive (SRA) and the European Nucleotide Archive (ENA) (Fig. 1), along with associated metadata. In order to ensure that MtExpress data are reliable and well documented, only sequencing data associated with peer-reviewed publications were conserved, representing 431 different samples (1374 if considering biological replicates). The nfcore/rnaseq bioinformatic pipeline version 3.0 (Patel et al. 2020) was used to automatically analyze the RNAseq libraries. First, FastQC/MultiQC were used to perform quality checking and reporting of sequencing data. Sequencing reads were, then, mapped to the Medicago genome version 5.1.7 (Pecrix et al. 2018) (52,723 genes) and reference genomes of Medicago symbionts or pathogens used in the selected SRA projects (listed in supplemental data set 1), using STAR mapper (Dobin et al. 2013). Finally, SALMON (Patro et al. 2017) was used to quantify read counts of transcripts. To enable cross-comparisons between different experiments, we normalized the complete dataset by the trimmed mean of M-values method (TMM) (Robinson and Oshlack 2010), using the EdgeR package. All samples were then manually labelled using a coherent nomenclature and organized into different categories based on organs or physiological conditions (18 categories in this first release; e.g. leaf, seed, abiotic factors, biotic stress, root nodule symbiosis…), with keywords attributed to each experiment, thereby enabling rapid search/selection.

**Figure 1.**
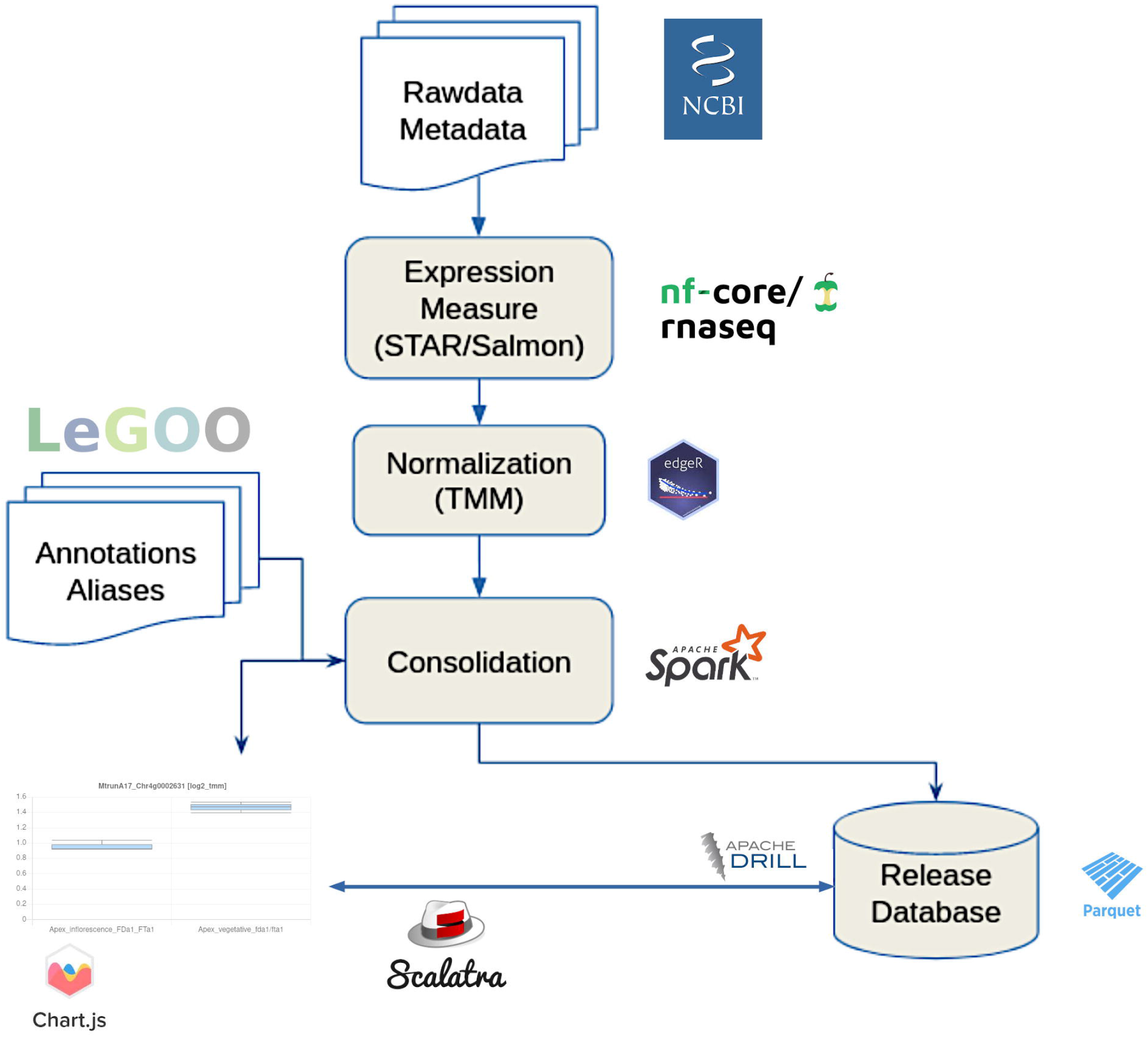
RNAseq analysis pipeline used to build MtExpress.

### A flexible and user-friendly interface

Thanks to synonymy links previously computed in the LeGOO knowledge base (Carrère et al. 2019), MtExpress can be queried by any gene locus identifier from the different *M. truncatula* genome releases (from Mt3.0 to Mt5.0), and identifiers of several microarray generations, namely Oli16k (Kuster et al. 2004), Affymetrix (Benedito et al. 2008) and Nimblegen (Verdier et al. 2013). Moreover, we extended this query flexibility to gene acronyms, based on a list manually extracted from publications, available on the Mt5.0 genome browser https://medicago.toulouse.inrae.fr/MtrunA17r5.0-ANR/. Furthermore, gene IDs from *Arabidopsis thaliana* (*A.thaliana* Col-0 [Araport11]) (Cheng et al. 2017) and four major legumes, namely soybean (*Glycine max* Williams 82 [Phytozome.12]) (Schmutz et al. 2010)), *Lotus japonicus* (*L. japonicus* Miyakojima MG-20 [3.0] (Li et al. 2020)), common bean (*Phaseolus vulgaris* BAT93 [EnsemblPlants.38] (Schmutz et al. 2014)) and pea (*Pisum sativum* cv. ‘Caméor’ v1a (Kreplak et al. 2019), can be used to get the expression pattern of putative orthologues in *M. truncatula* (Carrère et al. 2019).

The query output (Fig. 2) was set up to display by default the expression profile of the queried gene using a ‘reference dataset’, containing 28 samples which provide an overview of gene expression in various plant organs as well as symbiotic interactions. Then, the user can switch to one of the 18 pre-arranged categories, each with a more extensive or specialized set of samples: e.g. ‘abiotic factors’ and ‘biotic stress’ categories with all RNAseq samples related to different stresses, or ‘seed’ and ‘nodule’ categories, dedicated to samples specific to these organs. If needed, the complete dataset containing the 431 samples can also be selected for visualization. Finally, custom made selections can also be defined by the user by selecting/unselecting specific experiments to visualize and export the desired graphical outputs. All of the information associated to each experiment is provided, including the MultiQC (Ewels et al. 2016) report, and the corresponding publication, metadata and raw counts. Gene expression levels (with a choice between log2 TMM or raw TMM values) are displayed as box plots, with various informative elements (notably project name from SRA and descriptive statistics from biological replicates such as mean, median, min, max, Q1, Q3), each project being distinguished by a different color. Additional information on queried genes is easily accessible via the ‘synonymous’ and ‘annotation’ buttons, as well as links to the Mt5.0 genome browser and the LeGOO knowledge database. Conversely, a link to MtExpress is provided for all annotated genes in the Mt5.0 genome browser.

**Figure 2.**
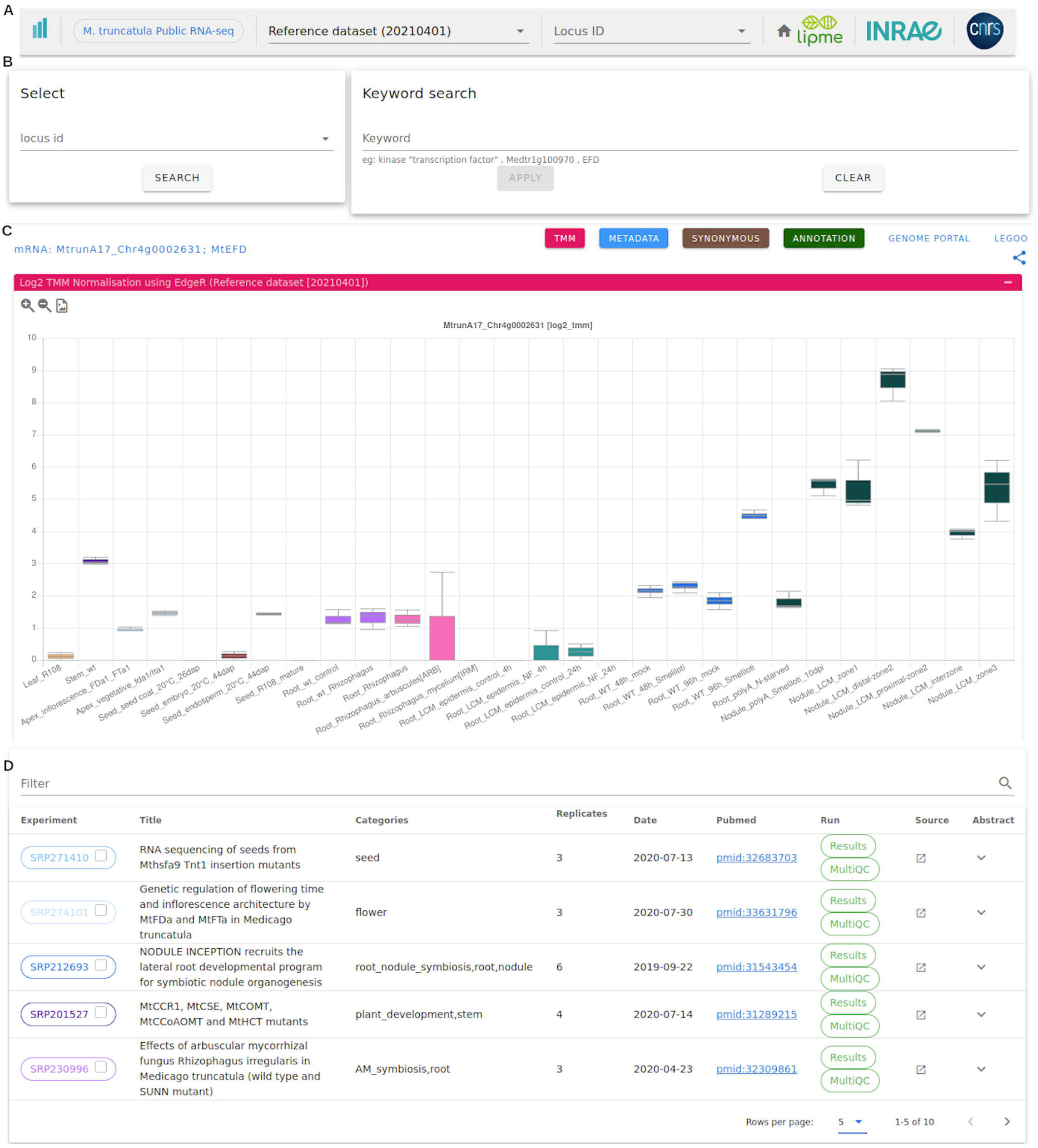
Interface of MtExpress, a user-friendly gene expression atlas. (A) drop-down menu for selecting dataset or sample category. (B) query boxes, using either Mt5.0 locus IDs, keywords, gene IDs from other genome releases or gene acronyms. (C) example of a graphical output using the reference dataset, with color-specific box plots per SRA project showing log2(TMM) values; buttons to switch between Raw and log2 TMM values and to display metadata, synonymous IDs and annotation information are indicated next to links to the Mt5.0 genome portal and the Legoo knowledge base. (D) list of plotted SRA projects with associated metadata and checkboxes to select/ unselect projects/samples.

To validate the overall pipeline, we examined the expression profile of a number of gene markers described to be specifically expressed in certain organs, tissues, or conditions. We found that their expression pattern was as expected from the literature. Three of them are shown in Fig. 3, using the reference sample set: MtPT4 (*PHOSPHATE TRANSPORTER 4*) (Harrison et al. 2002) (arbuscular mycorrhyzal symbiosis: arbuscule-containing cells), *MtNF-YA1 (NUCLEAR FACTOR YA1)* (Combier et al. 2006) (root nodule symbiosis: response to bacterial Nod factors, rhizobium infection, nodule inception and apical nodule zones), *MtDME (DEMETER)* (Satgé et al. 2016) (endosperm and nodule differentiation zone).

**Figure 3.**
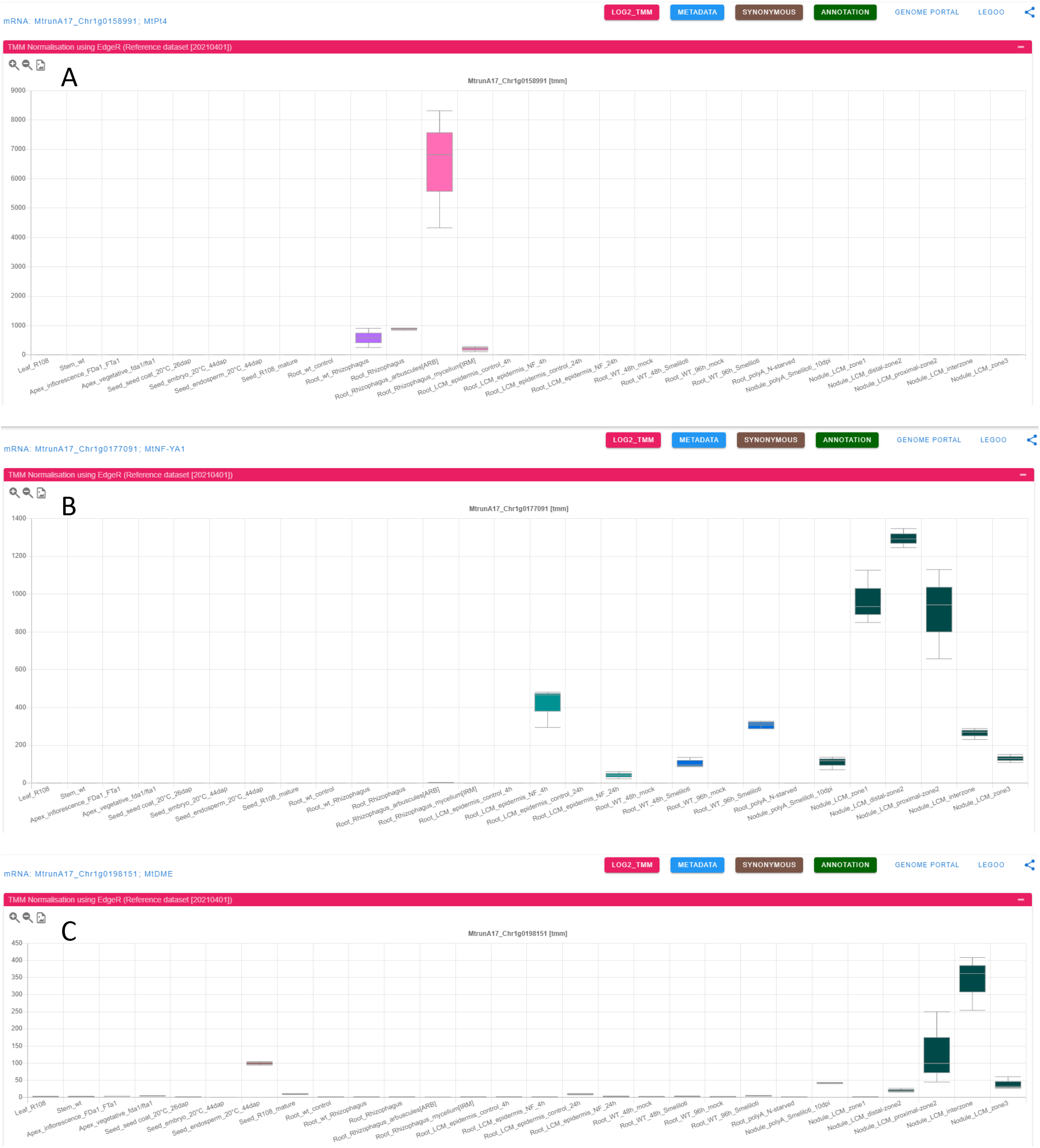
Expression profile of three genes with specific expression patterns. (A) MtPT4, expressed in arbuscule containing cells of mycorrhizal roots. (B) MtNF-YA1, expressed in bacterial Nod factor-treated epidermal root cells, in Rhizobium-inoculated root cells, and in nodule apical zones (zone 1, distal zone 2 and proximal zone 2). (C) MtDME, expressed in seed endosperm and in the nodule differentiation zones (proximal zone and interzone).

### Retrieval and presentation of MtGEA microarray data

As an additional tool to MtExpress, we downloaded publicly available Medicago Affymetrix gene chip data, previously stored in the Medicago Gene Expression Atlas (MtGEA) site hosted by the Noble Research Institute at https://mtgea.noble.org/v3. We implemented these data into the MtExpress webserver to provide an alternative solution to the inoperative MtGEA webserver. Moreover, we added features of MtExpress to this microarray-based gene expression atlas, such as the flexible input queries and pre-defined or custom-made categories. This alternative resource is also available at https://medicago.toulouse.inrae.fr/MtExpress.

## Discussion

MtExpress benefits from several important assets: (i) the use of recent releases of genome sequences/annotations to provide an up-to-date tool; (ii) semi-automated analysis pipelines allowing frequent updates (several times a year); (iii) manual re-labelling and organization of samples to conserve an unified structure through the updates; (iv) an attractive and flexible interface; (v) a transcriptomic platform fully integrated with other Medicago tools such as the Medicago Mt5.0 genome browser (Pecrix et al. 2018) and the LeGOO knowledge base (Carrère et al. 2019). The combination of manual curation with automated quality control and statistical analyses is key to ensure the reliability of MtExpress. Furthermore, the immediate access to accompanying metadata and publications represent an obvious gain of time and added value for users to better evaluate the data quality.

We therefore believe that this user-friendly, exhaustive and integrated resource will be of great use to the Medicago and, widely, to the legume community to quickly obtain the expression profiles of genes of interest in a variety of organs, tissues and conditions. This is, for instance, essential to define gene regulation patterns and determine gene expression specificity of developmental stage(s) or plant responses to biotic or abiotic environments. Considering the current pace of *M. truncatula* RNAseq data production and publication, we anticipate that the number and diversity of sequenced RNA samples will continue to grow rapidly. MtExpress will be updated accordingly, with creation of additional categories if needed. In the near future, we also plan to implement analytical tools such as gene co-expression analysis and pair-wise comparisons between experiments.

## Material and Methods

### Collecting Data

#### RNAseq dataset

We collected all data available as of April 1st 2021 in the Sequence Read Archive (SRA), using the Run Selector with the query “*Medicago truncatula* [ORG]” and the filter “AssayType = RNA-seq”. We kept all studies associated with a peer-reviewed publication, representing 51 projects altogether (supplemental Data set 1). All metadata coming from SRA, ENA and PubMed were associated to each selected project and stored for downstream treatment under a GIT repository [https://lipm-gitlab.toulouse.inra.fr/EXPRESSIONATLAS/ExpressionAtlasData]. Fastq files were directly downloaded from EBI SRA FTP mirror (ftp.sra.ebi.ac.uk) and listed in csv files with curated sample names, replicates and library strandedness to be properly processed.

#### Affymetrix microarray MtGEA dataset

We exported experiment replicates for all probesets (including Mtr, Msa, Sme and control probesets) from https://mtgea.noble.org/v3/experiments.php downloading webpage (Benedito et al. 2008). We excluded a few duplicated libraries and added one missing set of published data (supplemental Data set 1). All samples were manually ordered, classified into 18 categories (supplemental Data set 1) and aliases were added to homogenize labelling. Metadata such as publication, organ, experimental factors were manually extracted from the MtGEA website.

#### Expression Measure of RNAseq dataset

nf-core/rnaseq version 3.0 (Patel et al. 2020) was used with default parameters to run expression measure separately on each SRA project (Fig. 1). We chose STAR (Dobin et al. 2013) as read mapper and SALMON as read quantifier (Patro et al. 2017). We used the version 5.1.7 of *M. truncatula* A17 genome annotation (Pecrix et al. 2018) to map RNAseq data from different accessions of *M. truncatula*, as well as reference genomes of Medicago symbionts or pathogens used in the selected SRA Studies (*Sinorhizobium meliloti* 2011, *Sinorhizobium medicae* WSM419, *Fusarium oxysporum* f. sp. medicaginis, *Rhizophagus irregularis*, *Ralstonia pseudosolanacearum* GMI1000, *Verticillium albo-atrum* V31-2, *Gigaspora rosea* and *Rhizoctonia solani* AG-3) (supplemental Data set 1). Thanks to quality control reports and preliminary statistic analyses summarized using MultiQC (Ewels et al. 2016), we defined strandedness for each library and verified the consistency of biological replicates.

#### Normalization

##### RNAseq dataset

The complete dataset (51 SRA projects representing 431 samples) was normalized together in order to provide consistent values across gene expression comparisons in sub-categories, using TMM normalization from EdgeR package (Robinson and Oshlack 2010).

##### Affymetrix microarray MtGEA dataset

The expression data downloaded from the MtGEA site website were already normalized using a robust multichip average method (RMA normalization).

#### Consolidation

For RNAseq and microarray datasets, we added functional annotations provided by an automatic pipeline (Pecrix et al. 2018) as well as synonymy links to identifiers of different generations of *M. truncatula* microarrays and genome sequences/annotations (Carrère et al. 2019). We also used a manually curated list of 3,760 gene names/acronyms and products (Gene acronyms and IDs, release 20210409; https://medicago.toulouse.inra.fr/MtrunA17r5.0-ANR/). Normalized expression values (TMM or RMA) from biological replicates were summarized using descriptive statistics (mean, median, min, max, Q1, Q3) using adhoc Scala/Spark scripts to ensure future scalability. All these information were merged with dataset metadata to build a consistent database.

#### Database

The database consists of a repository of JSON and PARQUET files. Each dataset is documented through a JSON definition file along with run definition files including information related to reference genome and tools versions used for mapping. Database is accessed using Drill software (https://drill.apache.org/) version 1.16.0. Database structure and scheme are available at https://lipm-gitlab.toulouse.inra.fr/EXPRESSION-ATLAS

#### API

All normalized data are accessible through a REST API developed using the Scalatra framework and exposed with Jetty servlet container. The API is fully documented using Swagger 2.0. Documentation can be found at: https://lipm-browsers.toulouse.inra.fr/expression-atlas-api

#### User Interface

The user interface has been developed using the Vue.js library (v. 2.6.12) with a set of components, and plugins (see supplemental data set 1), notably Chart.js for boxplot display. It aims at providing simple functionalities such as gene searching through different accession identifiers (based on synonymous computed across different generations of genome assemblies/annotations and arrays), gene names or functions. Expression patterns are displayed using box-plots to show the level of variability between biological replicates. Users can easily select/unselect SRA projects to be plotted, navigate through different subsets of projects or locus and display raw or log2 normalized expression values, access to gene annotations, synonymous and links to relevant external resources (genome browser, knowledge base, original data repository). For each SRA project, links to MultiQC (Ewels et al. 2016) report, raw count and metadata files are provided.

#### Availability

Project name: MtExpress
Project home page: https://medicago.toulouse.inrae.fr/MtExpress
Source Code and Data: https://lipm-gitlab.toulouse.inra.fr/EXPRESSION-ATLAS
Operating system(s): Platform independent
Programming languages: Scala, R, Spark, Javascript, Perl
Other requirements: Drill 1.16, Apache web server
License: Apache License 2.0

## Supporting information

supplemental data set 1

## Acknowledgments

We thank Jérôme Gouzy and Marie-Françoise Jardinaud (LIPME, Castanet-Tolosan) for valuable discussions on the MtExpress set up and Nathalie Vialaneix (MIAT, Castanet-Tolosan) for her advice on statistical analyses. We also thank Erika Sallet, Ludovic Cottret, Ludovic Legrand and Léo Géré (LIPME) for technical advice. Finally, we are grateful to the Genotoul bioinformatics platform Toulouse Occitanie (Bioinfo Genotoul, https://doi.org/10.15454/1.5572369328961167E12) for providing computing resources.

